# Single-neuron bursts encode pathological oscillations in Parkinson’s disease and essential tremor

**DOI:** 10.1101/2022.04.05.486956

**Authors:** Maximilian Scherer, Leon A Steiner, Suneil K Kalia, Mojgan Hodaie, Andrea A Kühn, Andres M Lozano, William D Hutchison, Luka Milosevic

## Abstract

Deep brain stimulation procedures offer an invaluable opportunity to study disease through intracranial recordings from awake patients. Herein, we address the relationship between single-neuron and aggregate-level (local field potential; LFP) activities in the subthalamic nucleus (STN) and thalamic ventral intermediate nucleus (Vim) of patients with Parkinson’s disease (n=19) and essential tremor (n=16), respectively. Both disorders have been characterized by pathologically elevated LFP oscillations, as well as an increased tendency for neuronal bursting. Our findings suggest that periodic single-neuron bursts encode both pathophysiological beta (13-33Hz; STN) and tremor (4-10Hz; Vim) LFP oscillations, evidenced by strong time-frequency and phase-coupling relationships between the bursting and LFP signals. Spiking activity occurring outside of bursts had no relationship to the LFP. In STN, bursting activity most commonly preceded the LFP oscillation, suggesting that neuronal bursting generated within STN may give rise to an aggregate-level LFP oscillation. In Vim, LFP oscillations most commonly preceded bursting activity, suggesting that neuronal firing may be entrained by periodic afferent inputs. In both STN and Vim, the phase-coupling relationship between LFP and high-frequency oscillation (HFO) signals closely resembled the relationships between the LFP and single-neuron bursting. This suggests that periodic single-neuron bursting is likely representative of a higher spatial and temporal resolution readout of periodic increases in the amplitude of HFOs, which themselves may be a higher resolution readout of aggregate-level LFP oscillations. Overall, our results may reconcile “rate” and “oscillation” models of Parkinson’s disease and shed light onto the single-neuron basis and origin of pathophysiological oscillations in movement disorders.

**Significance:** In surgical patients with Parkinson’s disease and essential tremor, we leverage intracranial recordings to establish a link between pathophysiological phenomena across various scales of observation (spatio-temporal resolutions). We provide insights and reconcile theories about aberrant neurocircuit phenomena which underly theses debilitating, medically refractory movement disorders. Furthermore, our connectivity analyses between single-neuron and local field potential activities may shed light on the origin of the deleterious neural oscillations underlying these disorders. Ultimately, our findings may aid in the development or investigation of targeted therapies to address or correct underlying neurocircuit dysfunction, which can include neuropharmaceuticals, but also novel neuromodulatory strategies like closed-loop deep brain stimulation targeting pathophysiological oscillations and phase-dependent stimulation methods seeking to stimulate “at the right time/phase.”

## Introduction

The symptoms of Parkinson’s disease (akinetic-rigid features) and essential tremor are associated with pathologically elevated local field potential (LFP) activity in the subthalamic nucleus (STN; beta frequency oscillations; 13-33Hz) and the thalamic ventral intermediate nucleus (Vim; tremor frequency oscillations; 4-10Hz), respectively.^1,2^ Intraoperative microelectrode recordings enable acquisition of extracellular signals from these structures at both the single-neuron and aggregate neuronal (LFP) levels. LFPs are generally perceived as the conglomerate activity of individual neurons, with contributions from action potentials and subthreshold synaptic currents.^3^ In Parkinson’s disease, somatic spiking tends to cluster to specific phases of the beta LFP cycle,^4,5^ while such studies have not been performed for tremor-related oscillations in the Vim.

While beta frequency LFPs within the dorsolateral STN have been shown to correlate with clinical symptoms,^6^ evidence from animal models of Parkinson’s disease has also suggested that pathological phenotypes can be dissociated from beta oscillations.^7,8^ Instead, it was shown that functional impairment can be exclusively controlled by burst-firing patterns of the STN.^7^ Thus, the relationship between the STN LFP and somatic firing remains contentious.^8^ At the single-neuron resolution, STN neurons exhibit increased firing rates and a strong tendency to fire in bursts.^9,10^ Most recently, STN neurons that display burst firing patterns have been shown to represent a distinct, parvalbumin-positive neuronal population that, much like the source of STN beta frequency oscillations, cluster in the dorsolateral STN.^11,12^ Despite this topographical overlap, the relationship between neuronal bursting^13^ and LFP oscillations has not yet been characterized.

Electrophysiological studies have moreover demonstrated tremor-related LFP clusters in Vim,^14^ as well tremor-related single-neuron bursting that is congruent with peripheral tremor.^2,15^ However, mechanisms underlying the emergence of pathophysiological tremor-frequency oscillations have been elusive. Post-mortem studies have described various levels of cerebellar degeneration in patients with essential tremor,^16^ including lower levels of GABAergic tone.^17^ While lower inhibitory tone may contribute to increased disinhibition of deep cerebellar neurons, it does not explain the emergence of a periodic oscillation. However, a recent optogenetics study has suggested that synaptic pruning deficits of climbing fiber (CF) to Purkinje cell (PC) synapses may give rise to an increased propensity for cerebellar oscillations,^18^ which may subsequently spread throughout the cerebello-thalamo-cortical circuit.

In this work, we address the relationship between LFP and single-neuron activity in the STN and Vim of surgical patients. As *neuronal* (i.e., action potential) bursting^19^ appears to represent a pathophysiological hallmark in both Parkinson’s disease and essential tremor, single-neuron activity was separated into bursting and non-bursting episodes throughout the analyses. We moreover examined the temporal sequence between bursting and LFP activity using connectivity analyses to investigate whether periodic bursting may give rise to aggregate-level (LFP) oscillations, or whether the oscillation may in fact entrain neuronal firing. Ultimately, we aimed to provide insights as to whether the interaction between spiking activity and LFP conforms to a singular principle or whether this relationship needs to be investigated separately across different disease-relevant nodes and frequency bands.

## Materials and methods

### Data acquisition

Each patient underwent DBS implantation into the STN (akinetic-rigid dominant Parkinson’s disease; n=19) or VIM (essential tremor; n=16). During DBS surgery, intraoperative microelectrode recordings were used to localize STN^20^ or Vim.^21^ Single-neuron recording segments were extracted from 117 STN neurons and 32 Vim neurons; a data overview is provided in Supplementary Table 1. Acquired electrophysiological data were sampled at ≥12.5kHz using Guideline System GS3000 amplifiers (Axon Instruments, Union City, CA). Written informed consent was provided by all patients and the study was approved by the University Health Network Research Ethics Board.

For all analyses described below, the same methodological techniques were applied for STN and Vim recordings; however, analyses of STN recordings were focused on beta oscillations (13-33Hz), while analyses of Vim recordings were focused on tremor oscillations (4-10Hz).

### Extracting individual signals from the superimposed microelectrode data

Microelectrode recording signals were divided into several individual components. (1) LFP recordings were extracted using a lowpass FIR filter (50Hz) and (2) and a highpass FIR filter (300Hz) was used to better isolate spiking activity. Spiking activity was subsequently divided into subgroups of activity which occurred (3) during bursts and (4) outside of bursts (i.e., non-bursts). Burst detection was done via dissection of the inter-spike interval (ISI) probability distribution functions;^13,22^ Supplementary Fig. 1 contains details on ISI threshold setting for beta and tremor datasets. Undesired data points in signals (3) and (4) were masked by either zeros for subsequent power estimates or Gaussian noise for subsequent connectivity estimates.

### Spectral power in LFP and spiking signals

Power spectral density (PSD) estimates (i.e., Fig 1 and 3) were employed to determine the overall amplitude of beta/tremor activity for each of the delineated signals (1-4). While PSD estimations were directly calculated on the lowpass filtered LFP data (1), spiking activity signals (2-4) were first enveloped (absolute of the Hilbert transform). The PSD estimates were calculated using Welch’s^23^ method (1Hz bins, 50% overlap). PSD estimates obtained from unthresholded spiking signals (2) were used to classify each neuron as a strong or weak beta/tremor neuron in a semiautomated manner. An initial group of strong beta/tremor neurons were identified based on visual inspection of beta/tremor peaks in the PSD plots. The remaining neurons were subsequently classified using a naive Bayes classifier trained on the manually identified samples. For the relative height of the PSD peak at beta/tremor frequency to be automatically assessable, the absolute power at the peak frequency of interest was divided by an estimate derived from the interpolation of lower and higher frequency bins surrounding the peak.

**Figure 1:**
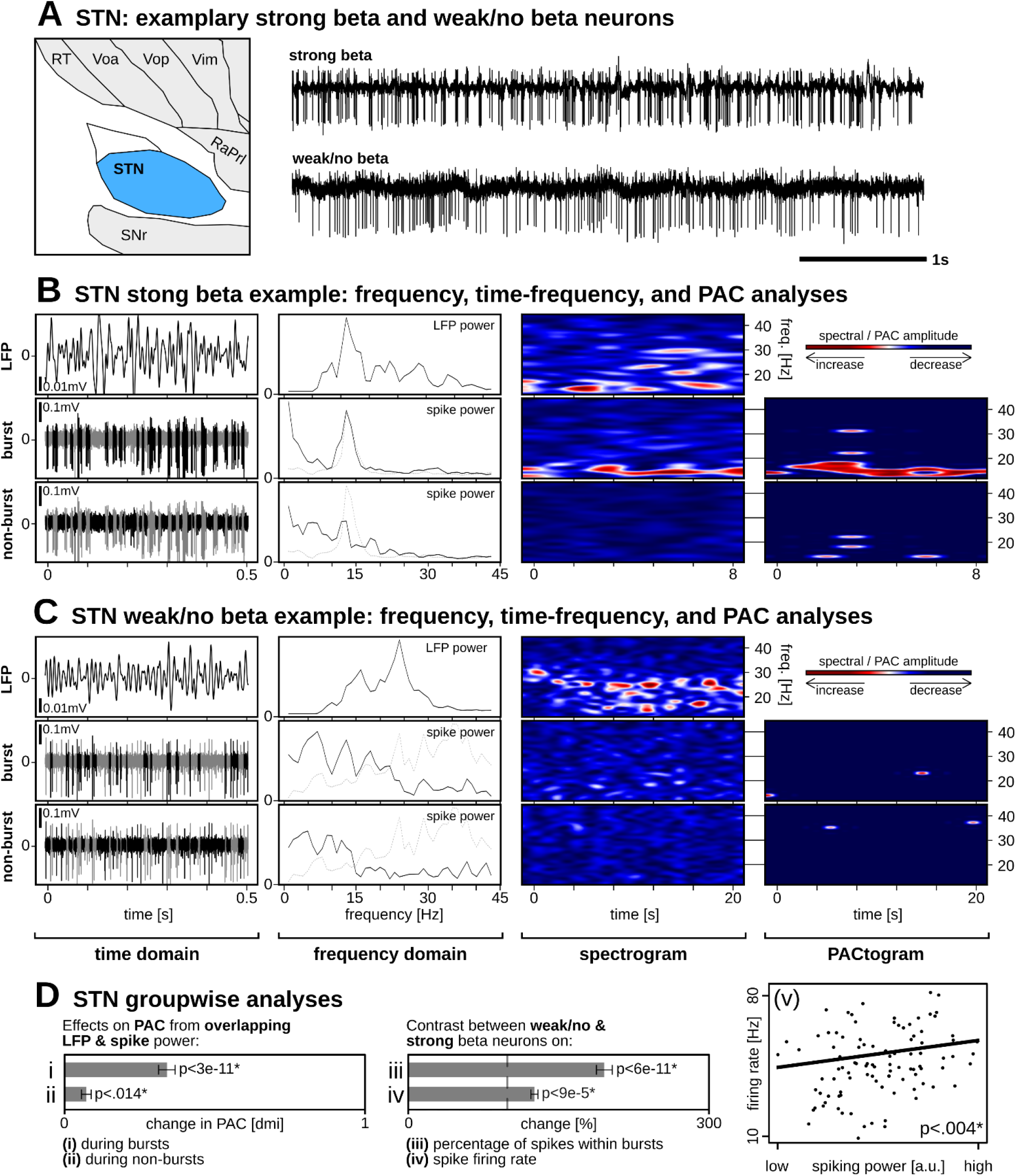
STN single-neuron and LFP activities. (A) Exemplary strong beta neuron showing periodic neuronal bursting and a weak/no beta neuron with minimal bursting. (B/C) Raw traces are divided into LFP, burst-thresholded, and non-burst-thresholded signal components. From left to right: same sample data as in (A) but at a higher time resolution; PSD estimates showing prominent frequency-matched peaks for LFP and burst-thresholded signals only; time-frequency spectrograms showing prominent frequency-matched oscillations for LFP and burst-thresholded signals only; PACtogram showing the strength of PAC over time between the LFP and corresponding single-neuron signals. (D) Groupwise statistics showing that (i) significantly increased PAC coincided with simultaneously increased LFP and spike PSD power during bursts, and (ii) during non-bursting activity although to a much lesser degree; (iii) the percentage of spikes within bursts and (iv) firing rates were significantly greater for strong compared to weak beta neurons; (v) there was a significant relationship between STN spike firing rate and PSD calculations performed on unthresholded spiking signal.

**Figure 2:**
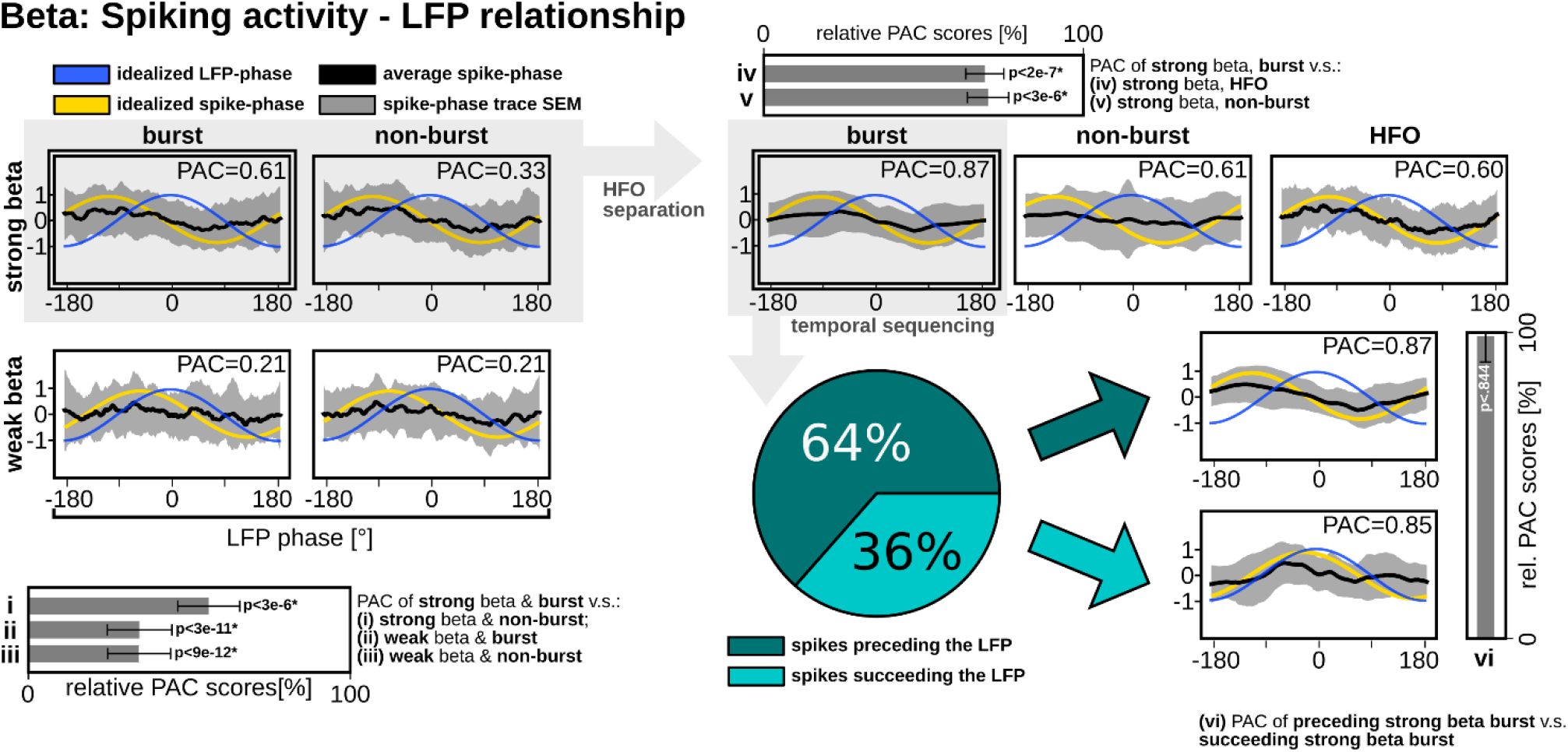
Groupwise phase-coupling between beta LFP and single-neuron activities in STN. (Left:) The greatest PAC relationship was found between LFP and strong beta burst-thresholded spiking and this relationship was significantly greater (i, ii, iii) than the three other conditions. (Top-right:) Upon removal of the HFO from strong beta spiking signals, the PAC sinusoidality of non-burst-thresholded spiking dissipated but was preserved for burst-thresholded spiking and the HFO only signal. PAC was significantly greater (iv, v) for burst-thresholded spiking than the other two conditions. (Bottom-right:) Based on SFC analyses, burst-thresholded spiking activity preceded the LFP oscillation 64% of the time; however, (vi) when signals were stratified by temporal sequence, PAC was not significantly different between the two conditions.

Time-frequency spectrograms were also generated from PSD estimates (beta: 2s windows, 1s window step size, and 1Hz wide bins; tremor: 4s windows, 0.25s window step size, and 0.25Hz wide bins). The “PACtograms” in Fig. 1 and Fig. 3 were generated from PAC estimates (described in detail in the following section) using 2s (beta) or 4s (tremor) data segments (beta: 1s window step size, and 1Hz wide bins; tremor: 0.25s window step size, and 0.25Hz wide bins) across various frequencies for the LFP’s phase component.

**Figure 3:**
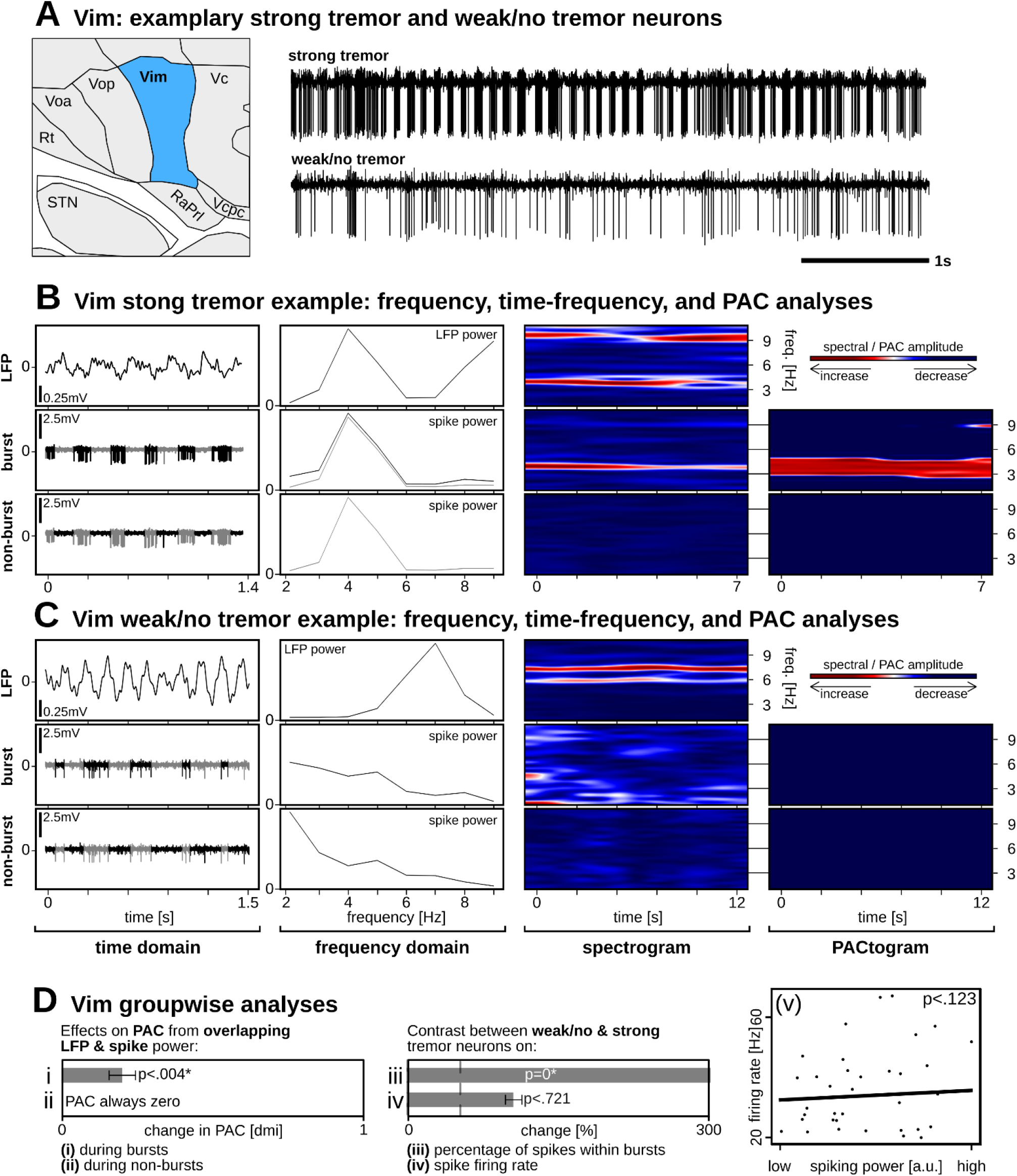
Vim single-neuron and LFP activities. (A) Exemplary strong tremor neuron showing periodic neuronal bursting and a weak/no tremor neuron with minimal bursting. (B/C) Raw traces are divided into LFP, burst-thresholded, and non-burst-thresholded signal components. From left to right: same sample data as in (A) but at a higher time resolution; PSD estimates showing prominent frequency-matched peaks for LFP and burst-thresholded signals only; time-frequency spectrograms showing prominent frequency-matched oscillations for LFP and burst-thresholded signals only; PACtogram showing the strength of PAC over time between the LFP and corresponding single-neuron signals. (D) Groupwise statistics showing that (i) significantly increased PAC coincided with simultaneously increased LFP and spike PSD power during bursts, but during non-burst spiking; (iii) the percentage of spikes within bursts was significantly greater for strong compared to weak tremor neurons, (iv) but no differences were found in firing rates; (v) nor was there a relationship between STN spike firing rate and PSD calculations.

### Connectivity analyses between LFP and spike signals

Phase-amplitude coupling (PAC) plots (Fig. 2 and Fig. 4) were generated by plotting the amplitude of spiking signals (3) or (4) with respect to the phase of each cycle of the corresponding LFP signal (1); positive values imply spiking activity whereas negative values imply lack of spiking. For the groupwise PAC histograms, amplitudes were normalized with respect to 25^th^ and 75^th^ percentiles on an individual sample level. A more sinusoidal and uniform PAC plot implies a stronger phase amplitude relationship between LFP and spiking activity. As such, an amplitude constrained (0.95-1.05) sinusoidal function was fit to each recorded sample individually. The resulting PAC score describes the goodness of fit between the fitted sinusoid and the sampled data points. The score was quantified as one minus the normalized mean squared error (MSE) between the sinusoidal fit and the sample LFP – spiking activity datapoints. The PAC plots also enabled the determination of the average phase preference between LFP and spiking activities. The phase offset was determined by comparing the groupwise fitted sinusoidal function (yellow curves in Fig. 2 and Fig. 4) to an idealized sinusoidal function with zero phase shift (blue curves in Fig. 2 and Fig. 4).

**Figure 4:**
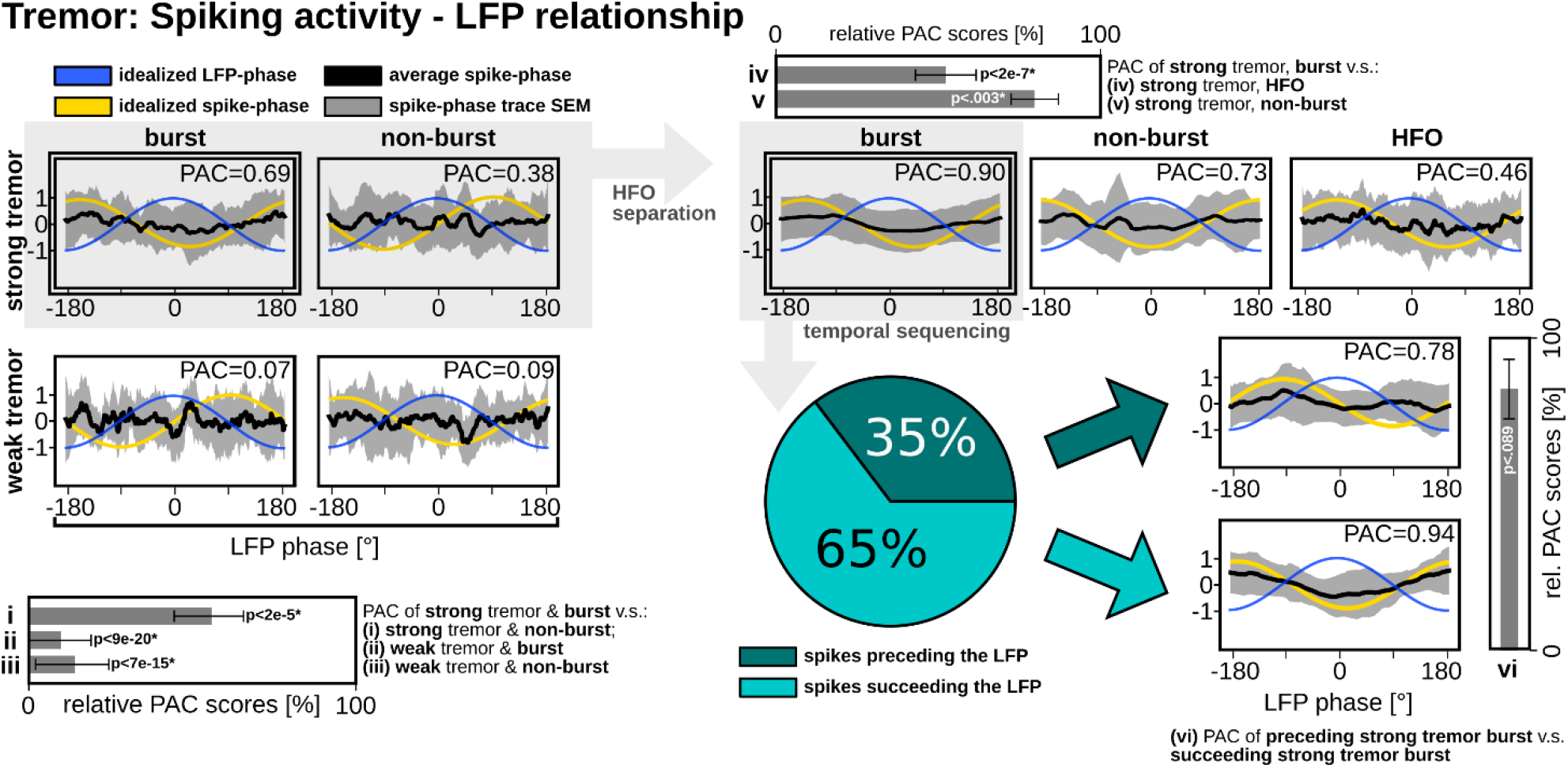
Groupwise phase-coupling between tremor LFP and single-neuron activities in Vim. (Left:) The greatest PAC relationship was found between LFP and strong beta burst-thresholded spiking and this relationship was significantly greater (i, ii, iii) than the three other conditions. (Top-right:) Removal of the HFO from strong beta spiking signals refined the sinusoidality of burst-thresholded spiking. PAC was significantly greater (iv, v) for burst-thresholded spiking than the other two conditions. (Bottom-right:) Based on SFC analyses, the LFP oscillation preceded burst-thresholded spiking activity 65% of the time; however, (vi) when signals were stratified by temporal sequence, PAC was not significantly different between the two conditions.

In a subsequent step, we took into consideration that each of signals (3) and (4) also contained high frequency activity (>300Hz) in addition to spiking activity. As only strong beta/tremor signals expressed high PAC values, the following analyses were exclusive to recordings from these neurons. Generally, high-frequency oscillations (HFOs), are thought to represent aggregate spiking activity, or neuronal noise. Previous reports have suggested that HFO amplitude modulation is a representation of synchronized spiking activity across a local cluster.^5^ Therefore, in subsequent analyses (Fig. 2 and Fig. 4), we removed HFO activity from the spiking signals (3) and (4) to generate: (5) burst-thersholded spiking with HFO removed, and (6) non-burst-thrsholded spiking with HFO removed. We also generated an (7) HFO only signal by removing spiking activity from signal (2). Unwanted data points were replaced with Gaussian white noise.

Temporal sequence between LFP and spiking activity (i.e., which signal precedes which) was determined using same frequency connectivity (SFC) estimates. SFC estimates were calculated from burst-thersholded spiking with HFO removed (5) of strong beta/tremor neurons as these signals expressed a much stronger PAC than their non-burst counterparts (6). SFC was estimated using directionalized absolute coherence (DAC; 3° minimal angle threshold). Briefly, DAC is a conglomerate SFC metric that quantifies direction of the information flow using the phase slope index, volume conductance via imaginary coherency, and magnitude of the relationship via magnitude squared coherence.^24^ Based on the results of SFC analysis, each individual recording was subdivided based on whether spiking activity preceded LFP activity or vice versa. PAC analyses were subsequently repeated on these subgroups (Fig. 2 and Fig. 4).

### Statistical analyses

Several groupwise statistical tests were computed based on the analyses presented in Fig. 1/2 for STN beta and Fig. 3/4 for Vim tremor. Firstly, we were interested in whether there were time and frequency dependent overlaps in the time-frequency spectrograms of LFP and spiking signals; and whether these overlaps in elevated spectral power coincided with the generated PACtogram plots. To investigate this, we used the time-frequency spectrograms from LFP and enveloped spiking activity to determine areas of above average activity. Then, we calculated the overlap between these areas across signals, and compared PAC within these areas to PAC outside of these areas using linear mixed models (LMMs) (Fig. 1Di/ii and Fig. 3Di/ii). For neuronal recordings, we also used LMMs to assess whether there was a difference in the percentage of spikes within bursts and the average firing rate when the recording sites were stratified by oscillatory strength (i.e., strong versus weak spectral power) (Fig. 1Diii/iv and Fig. 3Diii/iv). We also performed statistical tests using PAC scores seen in Fig. 2 and Fig. 4 by comparing each condition to the condition with the highest PAC score (i.e., the selected reference condition). For example, in Fig. 2A (left side); strong beta burst-thresholded spiking was used as a reference and PAC was compared to strong beta non-burst-thresholded, weak beta burst-thresholded, and weak beta non-burst-thresholded spiking. For all LMM evaluations, patient and trial IDs were included as random factors to compensate for multiple measurements across and within patients. Exact LMM models, parameters, and hypothesis counts for multiple comparison corrections (FDR corrected using the Benjamini-Hochberg method^25^) are available in Supplementary Tables 2 & 3.

### Data availability

Data are available within the Supplementary Material and Python code related to the analyses is available at https://github.com/Toronto-TNBS/pac_investigation.

## Results

### STN beta spectral power in LFP and spiking signals

PSD calculations were used to investigate beta peaks in the LFP signals, as seen in Fig. 1B/C. PSD calculations performed on enveloped spiking activity revealed that the major contributor to the PSD peak was neuronal bursting (e.g., Fig. 1B). Increased PAC coincided with simultaneously increased LFP and spiking activity power levels during bursts (p<6e-11*; Fig. 1Di), and during non-bursting activity (p<.014*; Fig. 1Dii) although to a much lesser degree. We were moreover interested to investigate if potential relationships existed between spike firing characteristics and PSD calculations. We found a greater percentage of spikes within bursts for strong compared to weak beta neurons (p<6e-11*; Fig. 1Diii), and that the average firing rate was greater for strong compared to weak beta neurons (p<9e-5*; Fig. 1Div). We moreover found a significant relationship between STN spike firing rate and PSD calculations performed on unthresholded enveloped spiking signal (p<.004*; Fig. 1Dv).

### STN beta connectivity between LFP and spike signals

Analyzing the PAC scores revealed initially that the highest PAC score was found for burst-thresholded spiking classified as having strong beta (Fig. 2 left side). In particular, the PAC scores of strong beta burst-thresholded spiking were higher than the PAC scores of strong beta non-burst-thresholded (p<3e-6*; Fig. 2i), weak beta burst-thresholded (p<3e-11*; Fig. 2ii), and weak beta non-burst-thresholded (p<9e-12*; Fig. 2iii) spiking. After separating HFO components (Fig. 2 right side, upper segment), strong beta burst-thresholded spiking with HFO removed showed a stronger PAC relationship than strong beta non-burst thresholded spiking with HFO removed (p<12e-7*; Fig 2iv) and HFO alone (p<3e-6*; Fig 2v). Notably, any semblance of PAC from non-burst-thresholded spiking was now abated upon removal of the HFO; but preserved within the HFO only signal.

Having identified that the most robust relationship between spiking and LFP activity was during periods of single-neuron bursting (when the HFO was removed), we subsequently analyzed the directional information flow between the strong beta burst-thresholded spiking with HFO removed and the LFP signal. This revealed that bursting activity preceded LFP activity in 64% of recordings, while bursting succeeded LFP activity in 36% of recordings (Fig. 2 bottom). We looked at PAC relationships once more after categorizing data by temporal sequence. PAC was marginally greater for bursts preceding the LFP, but not significant (p<.844; Fig. 2vi). Finally, preceding bursts expressed a clear sinusoidal distribution with a ~-110° phase preference.

### Vim tremor-band spectral power in LFP and spiking signals

As with STN beta, neuronal bursting was a major contributor to Vim tremor-related power spectrum (Fig. 3B). Increased PAC coincided with simultaneously increased LFP & spiking activity power levels only during bursts (p<.004*; Fig. 3Di), and not during non-bursting activity (where no PAC was present; thus, a statistic could not be calculated; Fig. 3Dii). We moreover found a greater percentage of spikes within bursts for strong compared to weak tremor neurons (p=0*; Fig. 3Diii) but no significant difference in firing rate between strong and weak tremor neurons (p<.721; Fig. 3Div). Moreover, we did not find a significant relationship between the Vim spike firing rate and PSD calculations performed on the unthresholded enveloped spiking power (p<.123; Fig. 3Dv).

### Vim tremor-band connectivity between LFP and spike signals

Analyzing the PAC scores revealed initially that the highest PAC score was found for strong tremor burst-thresholded spiking (Fig. 4 left side). In particular, PAC scores of strong tremor burst-thresholded spiking were higher than the PAC scores of strong tremor non-burst-thresholded (p<2e-5*; Fig. 4i), weak tremor burst-thresholded (p<9e-20*; Fig. 4ii), and weak tremor non-burst-thresholded (p<7e-15*; Fig. 4iii) spiking.

HFO separation revealed that strong beta burst-thresholded spiking with HFO removed was associated with a stronger PAC than strong beta non-burst thresholded spiking with HFO removed (p<2e-7*; Fig. 4iv) and HFO alone (p<0.003*; Fig. 4v).

SFC analyses revealed that strong beta burst-thresholded spiking with HFO removed preceded LFP activity in 35% of recordings, whereas bursting succeeded LFP activity in 65% of recordings (Fig. 4 bottom). This preferred temporal sequence was opposite to our observation for STN beta. We moreover found a small, but not statistically significant, (p<.089; Fig, 4vi) difference in PAC score when comparing neurons succeeding versus preceding LFP activity. Finally, succeeding bursts expressed a clear sinusoidal distribution with a ~±180° phase preference.

## Discussion

We explored the relationship between LFPs and single-neuron activity in the context of STN beta oscillations in Parkinson’s disease and Vim tremor oscillations in essential tremor. Discoveries of pathophysiological changes across various spatial and temporal resolutions have led to the development of mechanistic models to explain circuit dysfunction.^26^ In Parkinson’s disease for example, the “rate” model suggests that changes in single-neuron rates and patterns across nodes of the basal ganglia give rise to certain clinical symptoms of the disorder. The “oscillation” model suggests that local clusters of individual neurons synchronize their activity, giving rise to rhythmic, pathophysiologically-elevated aggregate-level oscillations. Although both models attempt to abstract pathophysiological processes of the same disorder, there has been little conceptual overlap, leading to the suggestion that these processes may be mutually exclusive.^7^ By exploring the relationship between single-neuron activity and aggregate-level LFP oscillations, we investigated whether rate and oscillatory neural correlates converge to a common electrophysiological signature or should be conceptualised as distinct hallmarks of the underlying pathophysiology. Overall, our results agree with previous studies which have suggested a relationship between spiking and LFP activity,^4,5,27^ but provide additional insights by identifying the critical role of neuronal bursting in facilitating this relationship across spatial and temporal resolutions.

### Neural coding with bursts to reconcile rate and oscillation theories

We found distinct peaks within power spectral estimations performed on enveloped spiking activity that corresponded to peaks within the power spectra of LFPs (Fig. 1 and Fig. 3). However, spiking activity thresholded to contain busting only resulted in far stronger power spectral estimates, whereas spiking activity occurring outside of bursts carried almost no oscillatory information. Moreover, we found a large degree of temporal overlap between the periodic bursting of single-neuron activity and the ongoing LFP oscillation; both confined to the same frequency band (Fig. 1Di and Fig. 3Di and corresponding PACtograms). These findings suggest that bursting activity within the spike train may represents a high spatial resolution representation of the aggregate level LFP oscillation (discussed in more detail in the following sections). Indeed, recordings with strong beta or strong tremor oscillatory activity in the LFP had more bursting than recordings with weaker oscillatory activity (Fig. 1Diii and Fig. 3Diii). Moreover, the percentage of spikes within bursts was greater in Vim recordings with strong tremor-related activity than in STN recordings with strong beta (p<2e-05*; not depicted), suggesting the transient^28,29^ nature of elevated beta LFPs compared to more sustained/enduring Vim tremor-related oscillations.

Neuronal bursting is a ubiquitous physiological means of neural information coding,^13^ serving several important functional roles in the brain. One such role is to increase reliability of information transfer across synapses.^30^ Pervasive neuronal bursting in STN and Vim may thereby represent a maladaptive change to brain circuitry, by which bursting activity pathologically exaggerates efferent information transfer. To this end, excessive STN and Vim bursting can indeed explain specific clinical features of Parkinson’s disease and essential tremor, respectively.

Our findings suggest important implications of neuronal bursting activity as a means of reconciling and corroborating the “rate”^31,32^ and “oscillation”^33^ models of Parkinson’s disease. The rate model suggests that dopaminergic degeneration (amongst other sequalae) results in decreased inhibition of the STN and subsequent over-excitation of the globus pallidus internus (GPi) and substantia nigra pars reticulata (SNr), which in turn over-inhibit thalamocortical and brainstem motor networks, resulting in the hypokinetic symptoms of Parkinson’s disease. In addition to increased firing rates, there is also an increased propensity for neuronal bursting within STN. Rather than considering the firing rates and patterns of individual neurons, the oscillation model implicates synchronous, aggregate-level network oscillations in the beta frequency band as a pathophysiological hallmark of the akinetic-rigid features of Parkinson’s disease.^6,34–36^ Within this work, we show a direct temporal link between neuronal bursting activity and the aggregate-level LFP oscillation (via PAC results); additionally corroborating the topographical overlap of these phenomena in STN.^11,12^ As such, we suggest that the rate model and oscillation model, both hallmarks of the hypokinetic features of Parkinson’s disease, are in fact *not* distinct and competing theories, but can be linked to one another through bursting. Our results suggest that increased beta activity in the STN LFP represents a functional readout of an increased propensity for periodic neuronal bursting (synchronized across many neurons), which is also associated with a net increase in neuronal output (Fig. 1Diii/iv). This explanation may therefore reconcile the rate and oscillation models of Parkinson’s disease (Fig. 1Dv), particularly given that both models implicate the hypokinetic features of the disorder and given that both firing rates and LFP oscillations are modulated by voluntary movements^37,38^ (Supplementary Fig. 2 & 3) and antiparkinsonian medications.^39,40^

A rate model in essential tremor has not previously been put forth; however, pathophysiologicaly elevated tremor-related neuronal bursting in the Vim is well-known phenomenon.^2^ Like with STN beta activity, we were able to establish a strong link between neuronal bursting and the tremor-frequency LFP signal in the Vim. Circuit dysfunction in essential tremor is indeed characterized by an increased propensity for periodic neuronal bursting across the cerebello-thalamo-cortical motor circuit, ultimately resulting in clinical/behavioural manifestations of tremor.^41^

### Connectivity between neural signals across spatial resolutions

We elucidated important temporal relationships between single-neuron bursting and the aggregate-level LFP oscillation. For STN, the highest degree of PAC between beta LFP and spiking activity occurred during bursting, particularly when the bursting activity preceded the LFP. Importantly, before removal of the HFO signal, a PAC relationship was nevertheless present between the beta LFP and non-burst signals within Fig. 3. Recent reports have indeed described a relationship between the phase of beta LFP signals and the amplitude of HFOs within the parkinsonian STN^5^ and motor cortex.^42^ However, removal of HFO components from the spiking signals completely abolished any semblance of a phase-amplitude relationship between the beta LFP and isolated nonburst spiking activity (Fig. 2 top right), while emphasizing the critical contributions of isolated single-neuron bursting activity to the beta LFP. As such, increased HFO amplitudes may indeed be driven by synchronous single-neuron bursting, and our results may thereby define and corroborate micro (single-neuron), meso (HFO), and macro (LFP) level representations of parkinsonian subthalamic beta oscillations across multiple scales of observation. The above results were also generally valid for tremor-related oscillations in the Vim, with three important differences: the PAC relationship was strongest when the tremor bursting in Vim *succeeded* LFP activity; the overall contribution of the HFO signal on non-bursting activity was less substantial; neuronal bursting phase preference was ±180° with respect to the LFP (compared to a −110° for STN).

The connectivity analysis of the temporal sequence between bursting and LFP may provide insights about the origin and propagation of pathophysiological oscillations within respective brain circuits and disorders. In particular, bursting activity preceding LFP oscillations (as was most common for STN beta) may suggest that coordinated synaptic events could underlie the emergence of bursting activity locally, which gives rise to an aggregate-level oscillation that may subsequently propagate throughout the circuit. On the other hand, LFP oscillations preceding spiking activity (as was most common for Vim tremor-related activity) may suggest that an incoming oscillation originating elsewhere (an external source) can entrain local neuronal activity, which can again continue to propagate throughout the circuit.

### On the emergence of STN beta oscillations in Parkinson’s disease

Exaggerated STN beta oscillations have indeed been correlated with clinical symptoms of Parkinson’s disease.^6,35,36^ At the initiation of elevated levels of LFP oscillations, coordinated action potential firing has been observed across cortex-basal ganglia structures.^4^ Critically, ensemble-level properties of synchronization have been shown to be underlain by the timing of action potentials in relation to cortical beta bursts, suggesting that cortical areas may be involved in the orchestration of beta activity through the basal-ganglia-thalamocortical loops.^4^ The importance of a cortical drive is further supported by recent evidence that has shown that the hyperdirect pathway is implicated in the transduction of beta oscillations to the STN.^43^ Repeated observations suggest that the synchrony between cortex and STN occurs predominantly at high beta frequencies (21-33Hz). However, it has been shown that low beta frequencies are more directly correlated with PD symptoms.^6,44–46^ Cortical control of the STN may still be implicated in the generation of this activity, as biophysical models have revealed that an exaggerated hyperdirect pathway can lead to the generation of subcortical synchrony at lower beta frequencies.^43^ This suggests that the dopamine depleted basal ganglia may provide a resonance circuit that amplifies cortical high beta, producing an exaggerated prevalence of pathological low frequency bursts. The maintenance of subthalamic beta oscillations has been suggested to be the result of the reciprocal connections of the STN with the GPe.^47^ In particular, prototypic GPe neurons have been shown to regulate beta activity by providing synaptic inhibition to STN neurons.^48^ When arriving in antiphase to a synchronized cortical drive,^49^ synaptic inhibition can support action potential generation by increasing the availability of Na+ channels, thus promoting the precision, efficiency, and ultimately the synchrony of STN spiking.^50^ Furthermore, the STN itself has been shown to be critical for the expression of beta oscillations in MPTP-treated monkeys.^51,52^

Overall, these studies implicate a role of coordinated synaptic mechanisms localized to the STN that are necessary for the generation and amplification of pathological beta oscillations. It is therefore possible that elevated LFP oscillations are only detectable when enough individual neurons are subject to these cascades of coordinated synaptic events which cause periodic oscillations of the firing rate (i.e., synchronized bursting); which would corroborate our findings that neuronal bursting most commonly precedes the LFP oscillation.

### On the emergence of Vim tremor oscillations

Essential tremor is regarded as a disorder of the cerebellum, evidenced by postmortem studies showing increased Purkinje cell loss^53^ and axonal swelling^54^ in the neocerebellum and vermis, as well as lower levels of GABA receptors in the dentate nucleus.^17^ However, these pathological changes do not directly explain the emergence of tremor-related oscillations. While unique ion channel dynamics in the thalamus, inferior olive, and cerebellum can generate oscillations via inhibition-induced rebound excitations,^55^ they have been reported to contribute to the generation of 10Hz physiological tremor only, but not essential tremor.^41^ Rather, there has been an important new theory that can directly explain the emergence of tremor-related oscillations based on essential tremor pathology. Recent studies in the postmortem cerebellum of patients with essential tremor have shown synaptic pruning deficits of CF to PC synapses.^56^ In particular, an increased number of CF synapses on the PC dendrites within the parallel fiber synaptic territory have been observed in essential tremor.^57^ These synaptic pruning deficits were linked to glutamate receptor delta 2 (GluRδ2) protein insufficiency.^18^ Critically, it was found that mice with GluRδ2 insufficiency and CF-PC synaptic pruning deficits develop essential tremor -like behaviour that could be suppressed with viral rescue of GluRδ2 protein.^18^ Moreover, optogenetic or pharmacological inhibition of neuronal firing, axonal activity, or synaptic vesicle release confirmed that excessive CF-to-PC synaptic activity was required for the generation of tremor and tremor-related oscillations in the cerebellum.^18^

As such, it is indeed likely that tremor-related oscillatory activity is generated within the cerebellum, which may subsequently spread to efferent structures, including the Vim. Strong periodic afferent inputs to Vim may therefore directly entrain local neuronal populations; which would corroborate our findings that neuronal bursting most commonly succeeds the LFP oscillation.

### Limitations

There are various limitations associated with intracranial neurophysiological studies in humans, including but not limited to time constraints and the inability to use pharmacological agents or sophisticated tools like optogenetics to elucidate ionic and molecular mechanisms of interest. On the other hand, these studies have an advantage in that it is not fully known how well animal models correspond to human conditions or anatomy. Moreover, while our connectivity results provided interesting insights regarding temporal sequence between signals, which may shed light on the origin of beta and tremor-related oscillations, the temporal sequences were not unanimous. The question of whether spiking activity drives the LFP or vice versa is in fact a ubiquitous one in neuroscience, which can only be unequivocally answered by simultaneous recording of far greater numbers of neurons than is currently feasible in humans or animals.^3^ Finally, it should be noted that a small proportion of STN neurons exhibited an antiphasic phase preference (70° versus the commonly observed −110°), which may be a result of differing levels of dopaminergic tone.^39^

## Conclusions

Periodic neuronal *bursting* likely encodes pathophysiological LFP oscillations in Parkinson’s disease and essential tremor, evidenced by strong PAC relationships between the two signals in the respective disorders. Synchronization between periodic neuronal bursting and the LFP oscillation may reconcile the canonical parkinsonian “rate” and “oscillation” models. Indeed, it was demonstrated that increased neuronal bursting was associated with elevated beta frequency oscillations. In STN, bursting activity most commonly preceded the LFP oscillation, suggesting that neuronal bursting generated within STN may give rise to an aggregate level LFP oscillation. In Vim, LFP oscillations most commonly preceded bursting activity, suggesting that neuronal firing may be entrained by periodic afferent inputs.

## Acknowledgements

The authors thank the patients for their participation.

## Funding

This work was supported by research grants from the University Health Network (L.M.), German Research Foundation (L.A.S., A.A.K.; Project-ID 424778381; TRR 295), Junior Clinician Scientist Program of the Berlin Institute of Health and German Academic Exchange Service (L.A.S.), and the Alexander von Humboldt Foundation (M.S.).

## Competing interests

Honoraria, travel funds, consultancy fees, and/or grant support unrelated to this work have been received from Medtronic (L.M., S.K.K., M.H., W.D.H, A.M.L.), Boston Scientific (S.K.K., A.M.L.), St. Jude-Abbott (A.M.L.), Inbrain Neuroelectronics (S.K.K.), Insightec (A.M.L.); none related to this work.

## Supplementary material

**Supplementary Table 1:** Data summary

**Supplementary Table 2:** Multiple comparison correction via the Benjamini-Hochberg method

**Supplementary Table 3:** Linear mixed models applied to evaluate the data

**Supplementary Figure 1:** Exemplary ISI distribution thresholding for burst detection

**Supplementary Figure 2:** STN modulation with movement

**Supplementary Figure 3:** Vim modulation with movement

## Notes

https://github.com/Toronto-TNBS/pac_investigation

