## supplementary_material for "Single-neuron bursts encode pathological oscillations in Parkinson’s disease and essential tremor"

### Contents

**Supplementary Table 1:** Patient table

**Supplementary Table 2:** Multiple comparison correction via the Benjamini-Hochberg method

**Supplementary Table 3:** Linear mixed models used to evaluate the data

**Supplementary Figure 1:** Exemplary ISI threshold estimation

**Supplementary Figure 2:** STN modulation with movement

**Supplementary Figure 3:** Vim modulation with movement

**Supplementary Table 1.** Data summary

| **Parkinson's disease** | | **Essential tremor** | |
|---|---|---|---|
| id | # neurons | Id | # neurons |
| 1 | 13 | 1 | 2 |
| 2 | 3 | 2 | 1 |
| 3 | 8 | 3 | 2 |
| 4 | 10 | 4 | 2 |
| 5 | 3 | 5 | 2 |
| 6 | 4 | 6 | 3 |
| 7 | 11 | 7 | 2 |
| 8 | 4 | 8 | 2 |
| 9 | 9 | 9 | 1 |
| 10 | 2 | 10 | 2 |
| 11 | 10 | 11 | 2 |
| 12 | 9 | 12 | 3 |
| 13 | 1 | 13 | 3 |
| 14 | 3 | 14 | 3 |
| 15 | 8 | 15 | 1 |
| 16 | 4 | 16 | 1 |
| 17 | 6 | | |
| 18 | 1 | | |
| 19 | 8 | | |

**Supplementary Table 2.** Multiple comparison correction via the Benjamini-Hochberg method

| significance | Benjamini-Hochberg value | k | p-values | - | data type | figure | feature |
|---|---|---|---|---|---|---|---|
| Yes | 0.00185185185185185 | 1 | 0 | | tremor | fig1 | burst |
| Yes | 0.0037037037037037 | 2 | 8.20E-20 | | tremor | fig4 | wb |
| Yes | 0.00555555555555556 | 3 | 6.19E-15 | | tremor | fig4 | wn |
| Yes | 0.00740740740740741 | 4 | 8.99E-12 | | beta | fig2 | wn |
| Yes | 0.0111111111111111 | 6 | 2.10E-11 | | beta | fig1 | area I |
| Yes | 0.00925925925925926 | 5 | 2.44E-11 | | beta | fig2 | wb |
| Yes | 0.012962962962963 | 7 | 5.42E-11 | | beta | fig1 | burst |
| Yes | 0.0148148148148148 | 8 | 2.54E-06 | | beta | fig2 | sn |
| Yes | 0.0166666666666667 | 9 | 1.23E-05 | | beta vs tremor | n/a | n/a |
| Yes | 0.0185185185185185 | 10 | 1.86E-05 | | tremor | fig4 | sn |
| Yes | 0.0203703703703704 | 11 | 8.46E-05 | | beta | fig1 | burst |
| Yes | 0.0222222222222222 | 12 | 0.00209873705171049 | | tremor | fig3 | corr – other |
| Yes | 0.0240740740740741 | 13 | 0.00351286167278886 | | beta | fig1 | corr – fig |
| Yes | 0.0259259259259259 | 14 | 1.31E-02 | | beta | fig1 | area II |
| Yes | 0.0277777777777778 | 15 | 0.0136278859680997 | | tremor | fig1 | area I |
| Yes | 0.0296296296296296 | 16 | 0.0267598442733288 | | beta | fig1 | corr – other |
| No | 0.0314814814814815 | 17 | 0.0393389463424683 | | tremor | fig3 | corr – other |
| No | 0.0333333333333333 | 18 | 0.0711065828800201 | | tremor | fig3 | corr – other |
| No | 0.0351851851851852 | 19 | 0.0717256665229797 | | tremor | fig3 | corr – other |
| No | 0.037037037037037 | 20 | 0.109199471771717 | | beta | fig1 | corr – other |
| No | 0.0388888888888889 | 21 | 0.12284317612648 | | tremor | fig3 | corr – fig |
| No | 0.0407407407407407 | 22 | 0.159568384289742 | | beta | fig1 | corr – other |
| No | 0.0425925925925926 | 23 | 0.27582249045372 | | beta | fig1 | corr – other |
| No | 0.0444444444444444 | 24 | 0.316422253847122 | | tremor | fig3 | corr – other |
| No | 0.0462962962962963 | 25 | 0.526523232460022 | | beta | fig1 | corr – other |
| No | 0.0481481481481482 | 26 | 0.720481059870868 | | tremor | fig1 | burst |
| | 0.05 | 27 | 1 | | tremor | fig1 | area II |
| alpha value | 0.05 | | | | | | |

**Supplementary Table 3.** Linear mixed models applied to evaluate the data

| **figure** | **data type** | **formula** |
|---|---|---|
| 1Di & 1Dii | beta | target_value ~ structured + (1\|patient_id) + (1\|trial) |
| 1Diii | beta | target_value ~ peak_type + (1\|patient) + (1\|trial) |
| 1Div | beta | target_value ~ peak_type + (1\|patient) + (1\|trial) |
| 1Dv | beta | target_value ~ hf_pow + (1\|patient_id) + (1\|trial) |
| 2i, 2ii, 2iii, 2iv, 2v, 2vi | beta | target_value ~ type + (1\|patient) + (1\|trial) |
| 3Di & 3Dii | tremor | target_value ~ structured + (1\|patient_id) + (1\|trial) |
| 3Diii | tremor | target_value ~ peak_type + (1\|patient) + (1\|trial) |
| 3Div | tremor | target_value ~ peak_type + (1\|patient) + (1\|trial) |
| 3Dv | tremor | target_value ~ hf_pow + (1\|patient_id) + (1\|trial) |
| 4i, 4ii, 4iii, 4iv, 4v, 4vi | tremor | target_value ~ type + (1\|patient) + (1\|trial) |

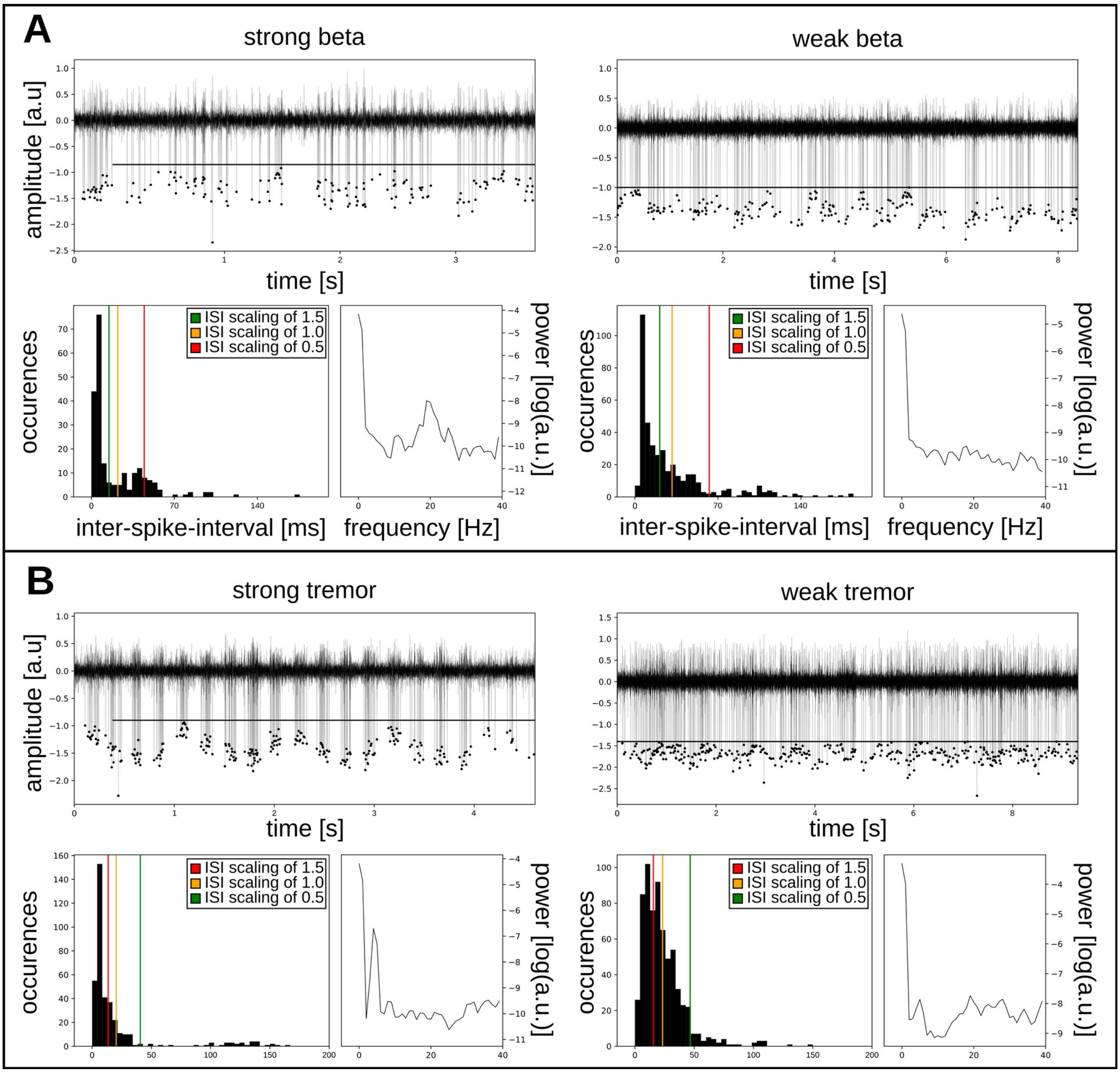

**Supplementary Figure 1: Exemplary ISI distribution thresholding for burst detection.** For strong beta neurons, two peaks were commonly discernable within the ISI distribution that could be accurately delineated at the bin corresponding to ~1.0 * the average ISI. For strong tremor, the two peaks could be accurately delineated at the bin corresponding to ~1.5 * the average ISI. These same thresholds were used for weak beta and tremor neurons, where two distinct peaks were not always as discernable.

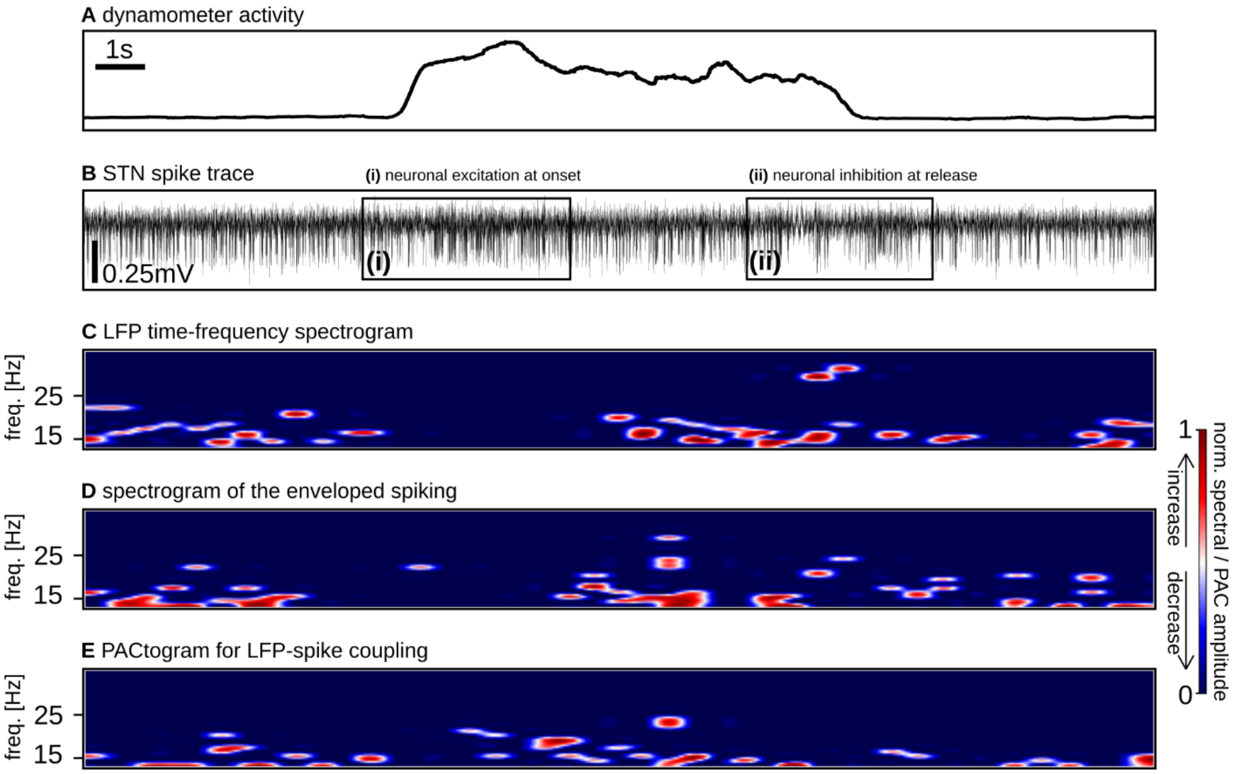

**Supplementary Figure 2: STN modulation with movement.** Exemplary data of STN neuronal, LFP, and LFP-spike coupling modulation during dynamometer squeeze. The disruption of beta-related neuronal bursting can be the result of (i) movement-related neuronal excitation or (ii) neuronal inhibition, both of which are associated with desynchronization of the LFP.

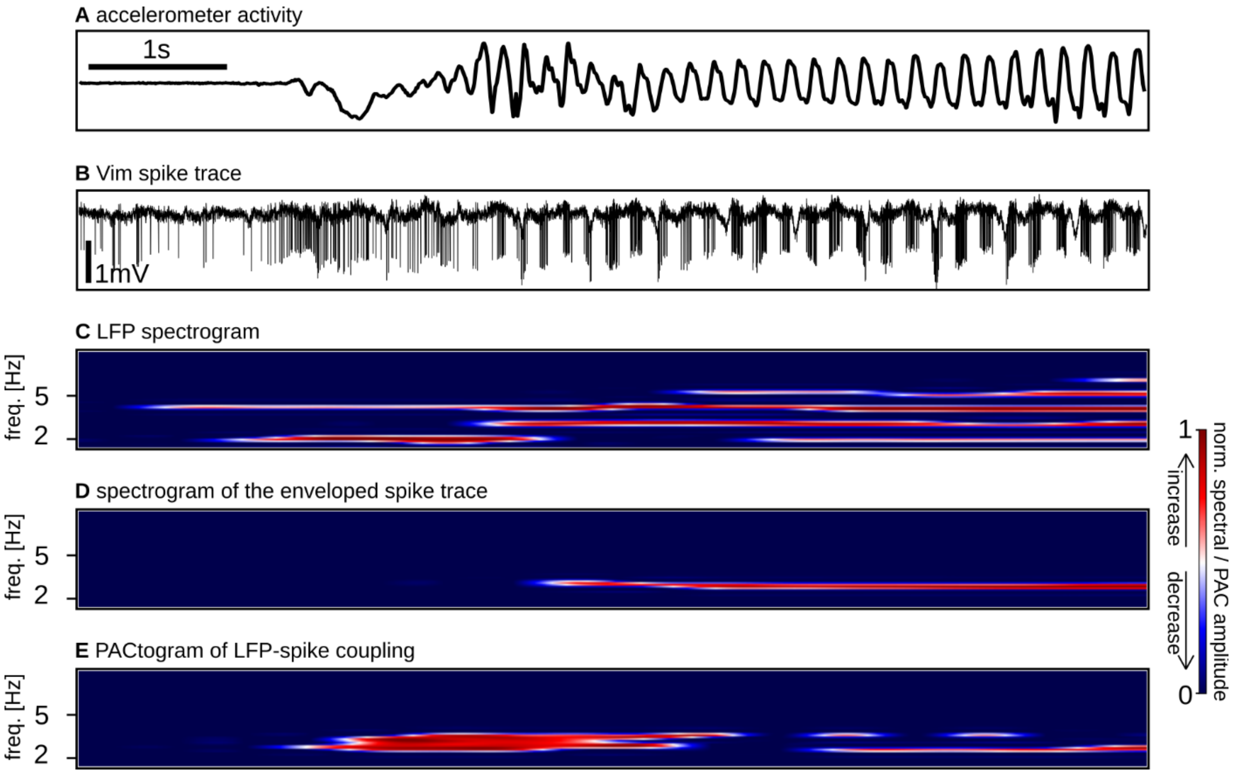

**Supplementary Figure 3: Vim modulation with movement.** Exemplary data of Vim neuronal, LFP, and LFP-spike coupling modulation during postural tremor onset measured using accelerometry (the patient moved their arm from a rest position to a postural hold which elicited their tremor). There is no tremor-related neuronal activity during rest, nor during the voluntary movement leading to the postural hold (during which there is a marked neuronal excitatory response).
